# A chronic photocapacitor implant for noninvasive neurostimulation with deep red light

**DOI:** 10.1101/2020.07.01.182113

**Authors:** Malin Silverå-Ejneby, Marie Jakešová, Jose J. Ferrero, Ludovico Migliaccio, Zifang Zhao, Magnus Berggren, Dion Khodagholy, Vedran Đerek, Jennifer Gelinas, Eric Daniel Głowacki

## Abstract

Implantable clinical neuroelectronic devices are limited by a lack of reliable, safe, and minimally invasive methods to wirelessly modulate neural tissue. Here, we address this challenge by using organic electrolytic photocapacitors (OEPCs) to perform chronic peripheral nerve stimulation via transduction of tissue-penetrating deep-red light into electrical signals. The operating principle of the OEPC relies on efficient charge generation by nanoscale organic semiconductors comprising nontoxic commercial pigments. OEPCs integrated on an ultrathin cuff are implanted, and light impulses at wavelengths in the tissue transparency window are used to stimulate from outside of the body. Typical stimulation parameters involve irradiation with pulses of 50-1000 μs length (638 or 660 nm), capable of actuating the implant about 10 mm below the skin. We detail how to benchmark performance parameters of OEPCs first *ex vivo*, and *in vivo* using a rat sciatic nerve. Incorporation of a microfabricated zip-tie mechanism enabled stable, long-term nerve implantation of OEPC devices in rats, with sustained ability to non-invasively mediate neurostimulation over 100 days. OEPC devices introduce a high performance, ultralow volume (0.1 mm^3^), biocompatible approach to wireless neuromodulation, with potential applicability to an array of clinical bioelectronics.

## Introduction

Implantable neural interfaces are at the heart of bioelectronic medicine, a growing field which aims to provide electrical solutions to medical problems^1–3^. Direct electrical actuation of the nervous system is utilized clinically in deep brain stimulation^4^, prosthetic retina implants^5^, vagus nerve stimulation for treatment of epilepsy^6^ and other disorders^7,8^, as well as numerous other applications. Meanwhile, the list of emerging technologies at a preclinical phase is constantly growing^9,10^. Several fundamental engineering hurdles need to be overcome to facilitate widespread implementation of bioelectronic devices and ensure optimal clinical outcomes^11,12^. A key challenge is to improve long-term powering and miniaturization of implantable devices, motivating exploration of methods to wirelessly actuate and control implants from outside of the body. The most common approaches involve radio frequency (RF) power transmission or electromagnetic induction^13^. Although these technologies are being developed for clinical use^11,14–16^, RF imposes size and shape constraints for transmitting and receiving components. The volume of the receiver (implanted inside the body) ranges from 30-600 mm^3^,^17^ including antenna for RF transmission, electrodes for nerve stimulation, and device packaging for protection of rigid Si-based electronics from body fluids. Efficient RF coupling and tissue heating are also factors that limit clinical translation.^17^ An alternative emerging approach leverages acoustic waves at ultrasound frequencies. Ultrasonic energy can be used to stimulate nervous tissue directly^18^ or can be absorbed by piezoelectric transducers to power devices^19^. Though very promising, due to acoustic impedance matching requirements the ultrasound transmitter must be in intimate contact with the skin, and penetration through layers of different tissues can be a limitation of ultrasound technologies. Overall, there is a strong demand for fully implantable systems which require less anatomical space than the aforementioned approaches, and which are as noninvasive and easy to use as possible.

In this work, we hypothesized that tissue-penetrating deep red light (620-800 nm) could effectively control implants wirelessly without requiring rigid or bulky implanted components. Three main features support this idea. Firstly, deep-red wavelengths occupy a “tissue transparency window” of the electromagnetic spectrum as they are neither absorbed by biopigments in the body nor by water. Therefore, unlike near-infrared radiation, they do not cause heating. Skin, muscle, and bone are all remarkably transparent to 620-800 nm light^20^. Next, light-emitting diode (LED) technologies are well established and mature. High brightness, efficient LEDs are reliable and commercially available in huge variety and at low-cost. Laser diodes give the additional advantage of low-divergence collimated beams that can target and deliver light through the tissue efficiently. Lastly, devices relying on optical power transfer can easily be made on the sub-millimeter scale (<1 mm^3^). Deep red light could therefore address the challenge of making small-scale devices that can be actuated and controlled wirelessly from outside the body.

The combination of an optoelectronic transducer implant and deep red light has been largely unexplored, with the notable exception of recent efforts to make light-powered/rechargeable pacemakers^21,22^. On the other hand, a variety of approaches in basic and biomedical research use light to mediate neurostimulation due to its noninvasiveness and versatility^23–25^. Optogenetics endows cells with light responsiveness^26^, but the necessity of genetic manipulation to accomplish this is not always facile or desirable for many applications, and remains a controversial proposition for clinical translation. Moreover, few opsins are available with light sensitivity in the red part of the spectrum^27^. These observations have spurred exploration of other ways to use light to interface with the nervous system over the past decade. Photothermal heating with light can be used directly to trigger a thermocapacitive effect, stimulating neurons *in vitro*^28^. To better control specificity and localization of this approach, light-absorbing nano or microparticles can be used as photothermal mediators^29^. Both inorganic^30^ (primarily silicon)^31^ and molecular^32^ or polymeric^33^ semiconductors can be used as light absorbers for various *in vitro* stimulation demonstrations. Nanoscale silicon biointerfaces can be tuned to provide highly-localized photocurrent stimulation in single cells^25^. Few of these concepts have proven scalable or reliable for *in vivo* settings, and chronic deployment remains elusive. To-date, photovoltaic neurostimulation has been developed for retinal protheses, based on arrays of silicon^34^ or organic semiconductors^35^. Highly optimized silicon diode-based technologies for retinal stimulation^36,37^ have advanced to clinical trials. Delivery of light is straightforward for intraocular applications. Getting sufficient light to devices implanted below skin and other tissues, however, is not as obvious and thus photovoltaic implanted neurostimulators have not been made. We propose that using organic semiconductors as the active optoelectronic component could facilitate light-mediated neurostimulation for such applications^38^ due to high absorbance coefficient, mechanical flexibility, and biocompatibility^39^. They can enable ultrathin and minimally invasive form factors inaccessible for traditional inorganic materials. Such approaches to noninvasive photostimulation of the nervous system *in vivo* have not yet been demonstrated.

Our biointerface devices use commercial organic pigments^32,40^, which belong to the category of safe and nontoxic colorants approved for a wide range of consumer products like food colorants and cosmetics^41^. These pigment materials form the basis of our recently developed organic electrolytic photocapacitor (OEPC)^42^. This device employs a nanocrystalline donor-acceptor PN junction acting as the charge-generating element and primary stimulation electrode, which is surrounded by a concentric return electrode. The stimulation efficacy of this minimalistic device architecture was validated for cultured neurons and explanted retinal tissues^42^. More recently, the capacitive charging behavior was characterized on the single-cell level, accompanied by direct electrophysiological measurement of the device’s impact on voltage-gated ion channels^43,44^. These *in vitro* demonstrations were based on rigid OEPCs. Here we integrate the OEPC into an ultrathin flexible architecture suitable for chronic *in vivo* implantation. The photoelectrical charging behavior of the OEPC stimulation devices effectively activated the rat sciatic nerve *in vivo* and enabled precise control of stimulation by varying light intensity and pulse duration. We fabricated the OEPC into a self-locking ultrathin cuff that was simple to surgically place and immobilize inside a freely-moving animal over months. Deep red light delivered through the skin surface to the implanted OEPC evoked compound muscle action potentials (CMAPs) via sciatic nerve stimulation at an operation depth of 10-15 mm. Device implantation did not impede physiologic motor behaviors, and devices maintained their operation for up to 3 months after implantation. With a total volume of 0.1 mm^3^, OEPCs are the lowest-volume wireless peripheral nerve interface reported to-date^17^. These results suggest that OEPCs provide a viable approach to chronic *in vivo* neurostimulation, and hold potential for translation to clinical applications.

## Results

### Design and fabrication of flexible OEPC nerve stimulators

Our approach to wireless neurostimulation leverages organic molecular thin-films to efficiently transduce light impulses into electrolytic currents that modulate activity of excitable cells. As in our *in vitro* studies leading up to the present work^42,43^, we rely on the phthalocyanine (H_2_Pc, P-type) / *N,N’*-dimethyl perylenetetracarboxylic bisimide / (PTCDI, N-type) heterojunction to create OEPCs (Figure 1a). The bilayer is deposited by thermal vacuum evaporation to form a densely-packed film of nanocrystals (Figure 1b). This PN junction combination has high reliability and stability in aqueous environments. To transform the OEPC into an implantable device capable of efficiently delivering stimulation current, we integrated it into a ribbon-like structure that can conform around the nerve. We used thin (5 μm) parylene C, a well-established biocompatible polymer, as a substrate material^45^. The first design challenge we faced was producing a semitransparent conducting back electrode layer on the substrate. A key materials selection criterion is semitransparency of this underlying conductor, to allow light to reach the absorbing PN semiconductor layer. We utilized thin thermally-evaporated Au (10 nm thickness) due to its excellent conductivity, good transparency, and mechanical flexibility^42^. In the OEPC architecture, the back conductor functions as a return electrode, providing a termination of the current path generated by the PN junction. The PN junction, upon illumination, will produce an electrolytic double layer as electrons accumulate at the N-type material/electrolyte interface. Meanwhile, holes are driven into the underlying metallic conductor, creating an oppositely-charged double layer around the PN pixel thus giving rise to ionic currents around the device (*I*_ionic_, Figure 1c). From the point of view of an underlying nerve, this device architecture produces a tripolar-type stimulating electrode arrangement (Figure 1c inset)^12^. These materials were integrated to ultimately fabricate chronically implantable photocapacitors (CIPs) for testing in an animal model. We placed these devices on rat sciatic nerve intra-operatively, and subsequently implanted them for long-term *in vivo* evaluation (Figure 1d). During chronic photostimulation, 638 nm light impulses would need to be beamed through about 10 mm of tissue to drive the CIP (Figure 1e). We first sought to investigate the feasibility of this approach – can light be transmitted efficiently to reach a device located below the surface of the skin? Light propagation through different tissues has been studied in detail^20^. We applied established numerical Monte Carlo (MC) methods^46,47^, and used known optical constants for rat tissue^48^ to determine that a conventional 700 mW laser diode at 638 nm will deliver light intensities in the range of tens of mW/cm^2^ at a depth of 1 cm (Figure 1f).

**Figure 1.**
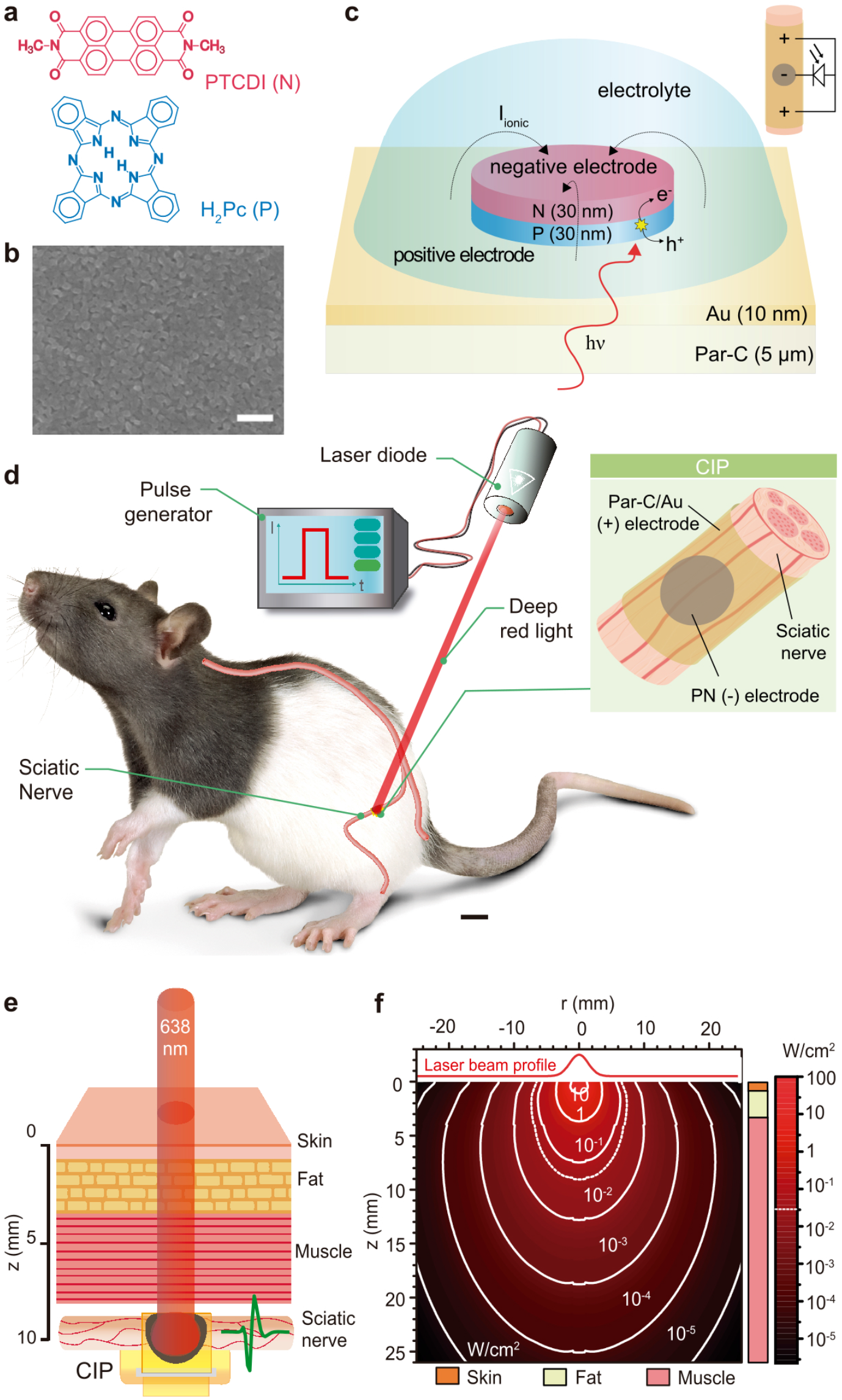
Organic electrolytic photocapacitors (OEPCs) wirelessly stimulate the sciatic nerve *in vivo*. **a,** Molecular structures of the active components in the PN semiconducting layer. Phthalocyanine (H_2_Pc) functions as the light-absorbing and electron-donating P-type layer. *N,N’*-dimethyl tetracarboxylic diimide (PTCDI) acts as the electron-accepting N-type layer and forms an electrolytic contact with the surrounding electrolyte. **b,** The sequentially evaporated PN bilayer (30+30 nm) forms a compact thin film featuring a distinctive nanocrystalline morphology apparent in scanning electron microscopy. Scale bar 200 nm. **c,** Diagram of the OEPC device and its operating principle. The PN bilayer is processed on top of a semitransparent gold film (10 nm), which acts as the return (+) electrolytic contact. Light in the deep-red region (638 or 660 nm) is used for excitation of the P-type layer. Photogenerated electrons travel through the N layer to accumulate at the electrolyte interface, forming an electrolytic double layer. Concurrently, holes are driven from the P layer into the underlying gold, forming an oppositely-charged double layer. During charging of the device (beginning of light pulse), and discharging, (end of light pulse), ionic displacement currents (transient ionic current, *I*_ionic_) flow around the device and thus through surrounding tissue. This produces biphasic stimulation pulses. The inset shows how the OEPC, understood simply as an illuminated photodiode, couples to a nerve with a quasi-tripolar arrangement. **d,** Schematic of the *in vivo* implanted OEPC photostimulation experiments performed in this study. Scale bar 1 cm. The inset details how a chronically-implantable photocapacitor (CIP) cuff is placed around the nerve, with the configuration of the primary PN photoelectrode (−) with the surrounding (+) return electrode. **e,** Following implantation, deep red light penetrates through skin, fat, and muscle tissues to reach the OEPC, located at a depth of roughly 10 mm. **f,** Numerical calculation of penetration of a stimulating red light beam through simulated animal tissue layers, showing the effectiveness of 638 nm light to access the implanted OEPC. The dotted white line represents the cross-sectional region with 50 mW/cm^2^ of intensity.

### Photoelectrical characterization

While stimulation electrode benchmarking methods are well-established^49^, determining relevant parameters for an electrically-floating photoelectrode device requires special consideration. A common figure of merit for a neurostimulation electrode is the electrolytic charge density that the electrode can inject, or charge density per phase^49^. In order to measure this parameter in the case of OEPCs, we devised the electrophotoresponse (EPR) method^43^ (Figure 2a). In EPR, the photovoltage or photocurrent is registered by contacting the back electrode (Au) with a probe and measuring versus an ideally nonpolarizable electrode (Ag/AgCl) immersed in an electrolyte. This electrolyte is confined to the top of the PN semiconductor region. The device is illuminated with light pulses from the bottom, through the semitransparent gold film, thus mimicking the anticipated configuration during *in vivo* neurostimulation. The photovoltage to which the PN/electrolyte junction charges can be measured using an oscilloscope. Photocurrent is quantified in the same arrangement using a current amplifier. During a light pulse of 638 or 660 nm, the OEPC charges to around 300-320 mV and the capacitive displacement currents under the same conditions peak at around 2 mA/cm^2^ (Figure 2b). Dynamics are rapid, with full charging voltage achieved within 20 μs. In the context of OEPC devices, charge density can be modulated with light intensity and light pulse length. We plot the charge density per phase as a function of light intensity (Figure 2c) and light pulse duration, from 50 μs to 1 ms (Figure 2d). The 638 nm wavelength was found to have slightly better photocharging efficiency relative to 660 nm. This finding corresponds to the absorption spectrum of the H_2_Pc p-type absorber layer^42^. Importantly, both wavelengths correspond to readily-available LED illuminators.

**Figure 2.**
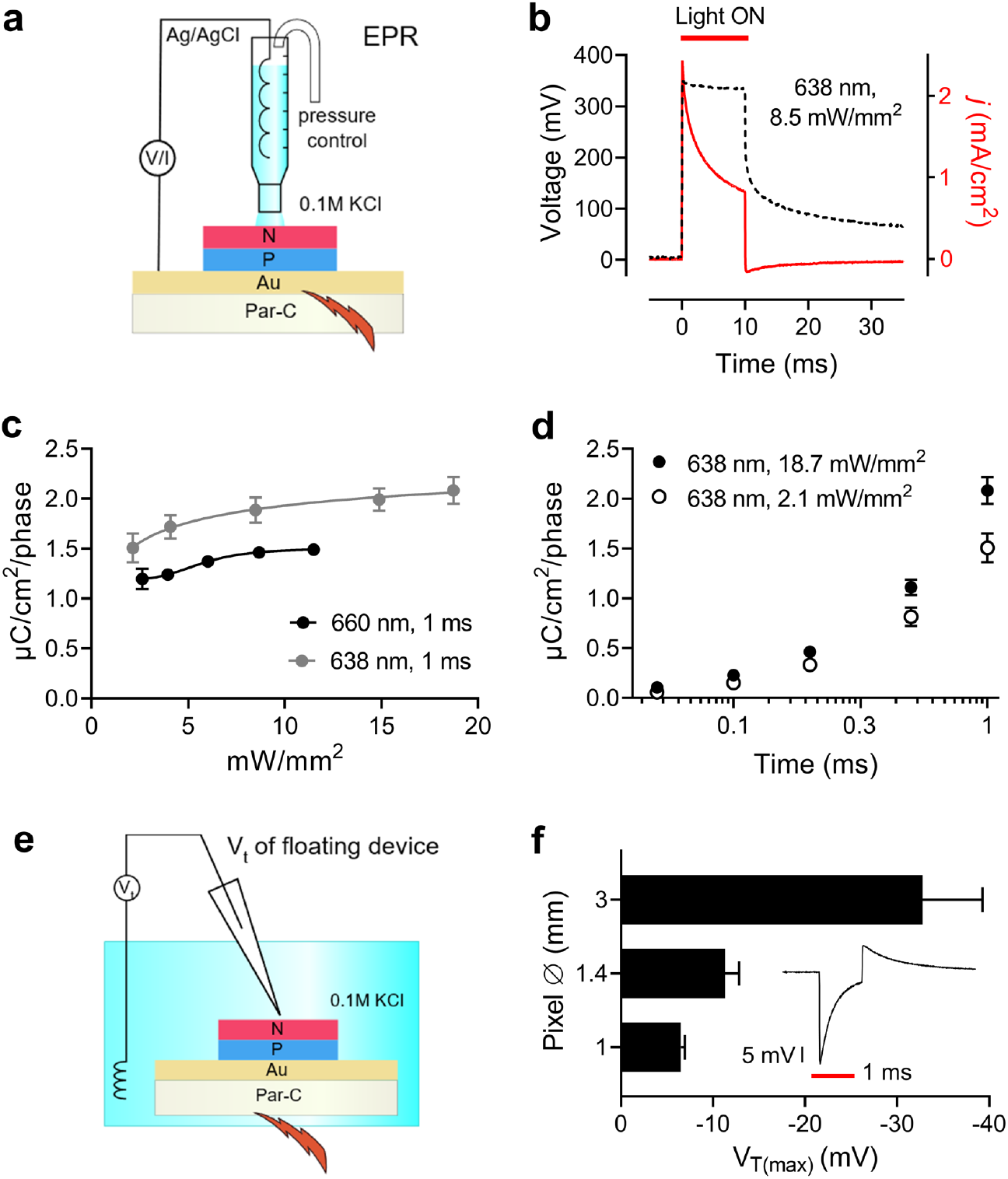
Photostimulated OEPC devices deliver rapid localized electrolytic pulses. **a,** Schematic of the electrophotoresponse (EPR) closed-circuit measurement, where photovoltage or photocurrent is measured between the bottom metal contact and an electrolytic contact. This method probes photogenerated charge injection across the PN/electrolyte junction. **b,** Photovoltage and photocurrent registered by EPR, using excitation with a 638 nm laser diode. Devices charge to full voltage within 20 μs. **c,** Charge density per phase as a function of light pulse intensity, for 1 ms light pulses of either 638 or 660 nm light, n=2, ±SD. **d,** Charge density per phase as a function of light pulse duration, n=2, ±SD. **e,** Transient voltage, V_T_, probes the voltage perturbation generated in electrolyte when displacement currents flow from anode (Au) to cathode (PN). V_T_ is registered between a point above the center of the PN pixel versus a distant Ag/AgCl reference. This method reflects the actual electrical perturbation nerve fibers would experience from the OEPC stimulator. **f,** V_T_ during a 1 ms light pulse, showing the clear biphasic current behavior. All pixels give qualitatively the same trace, only the magnitude of voltage varies with pixel diameter. The bar graph plots the cathodic maximum V_T_ as a function of PN pixel diameter. n = 3 for each size, ±SD.

While the EPR measurement allows quantification of photocharge density which the OEPC device can generate, it does not faithfully reflect the final operating conditions of an OEPC stimulator. *In vivo*, the OEPC device is electrically floating. The organic PN pixel forms an electrolytic closed circuit with the back electrode.^42,50^ The displacement current which flows around the device during photocharging and discharging generates transient potentials in the solution which affect the electrophysiology of nearby cells, as established in our earlier *in vitro* studies^43,44^. To visualize this effect, we measure the transient potential (V_T_) above the PN pixel in electrolyte (Figure 2e). The V_T_ was registered between a recording microelectrode positioned at 1 mm above the center of the PN pixel versus a distant reference electrode. Consistent with the PN polarity of the OEPC, illumination resulted in a cathodic transient voltage peak followed by an anodic transient when the light was turned off and electronic charge carriers recombine (Figure 2f inset). This voltage transient corresponds temporally to the electrical perturbation that adjacent axon bodies will experience. It mimics a charge-balanced biphasic stimulation protocol, which is typically used to avoid tissue and electrode damage during neurostimulation^51,52^. The actual transmembrane potential induced will vary based on the position and distance from the stimulating electrode^43,53^. The magnitude of the V_T_ was directly proportional to the size of the PN pixel, ranging from around 8 mV for a 1 mm diameter PN pixel to 30-40 mV for a 3 mm diameter pixel (Figure 2f). Taken together, these results demonstrate the operating mechanism and parameters for OEPC-mediated neural stimulation and highlight the importance of PN pixel size as a key parameter governing effectiveness of neuromodulation.

### Acute validation of light-induced nerve stimulation

To test the efficacy of *in vivo* neuromodulation using OEPC devices, we used the well-established rat sciatic nerve model^12,16^. To determine the effectiveness of OEPC-mediated nerve stimulation, microwires capable of measuring evoked compound muscle action potentials (CMAPs) were placed in biceps femoris (Figure 3a) and plantar muscle groups (Supplementary Figure S1) of 5 rats. OEPC devices integrated into parylene C ribbons were wrapped around the nerve with the PN pixel immediately adjacent to the epineurium. The ribbon was not fixed in any way other than simple adhesion of the loose plastic ends to each other via capillary interactions in the presence of water. Intra-operatively, OEPC ribbons with differently sized PN pixels (1 mm ø, 1.4 mm ø, and finally 3 mm ø, denoted henceforth as three device sizes S, M, and L) were illuminated using an LED (660 nm) with a collimator lens that was placed above the exposed nerve (Figure 3b). OEPCs were consistently placed such that the PN pixel pointed in the dorsocaudal direction relative to the rat’s body. To ensure that any recorded CMAPs were the result of light-mediated photovoltaic stimulation, we first placed a sham device (bare parylene C covered with Au in the absence of a PN pixel) on the nerve. Illumination of this sham device did not yield any response or artefact (Figure 3c, Supplementary Figure S1b). In contrast, illumination of OEPC devices stimulated the sciatic nerve, as demonstrated by visually observable twitching in sciatic-innervated muscles and recorded CMAPs (Figure 3d: CMAPs in response to 1-ms, 9.4 mW/mm^2^, illumination pulses; see also Supplementary Figure S1c, and Supplementary Video 1). Increased size of the PN pixel strongly correlated with higher stimulation as measured by larger amplitude of both observable movement and the average CMAP waveform (Figure 3d, Supplementary Figure S1c), concurring with the predictions of V_T_ measurements (Figure 2f). The elicited CMAPs were also highly consistent in both amplitude and latency after repetitive light-stimulation (Figure 3e, Figure S1d; 25 light-pules, 3 seconds between). Taken together, these results demonstrate the ability of the OEPC to transduce light impulses to electrical potentials that are capable of consistently and repetitively stimulating the sciatic nerve, with response magnitude dependent on PN pixel size.

**Figure 3.**
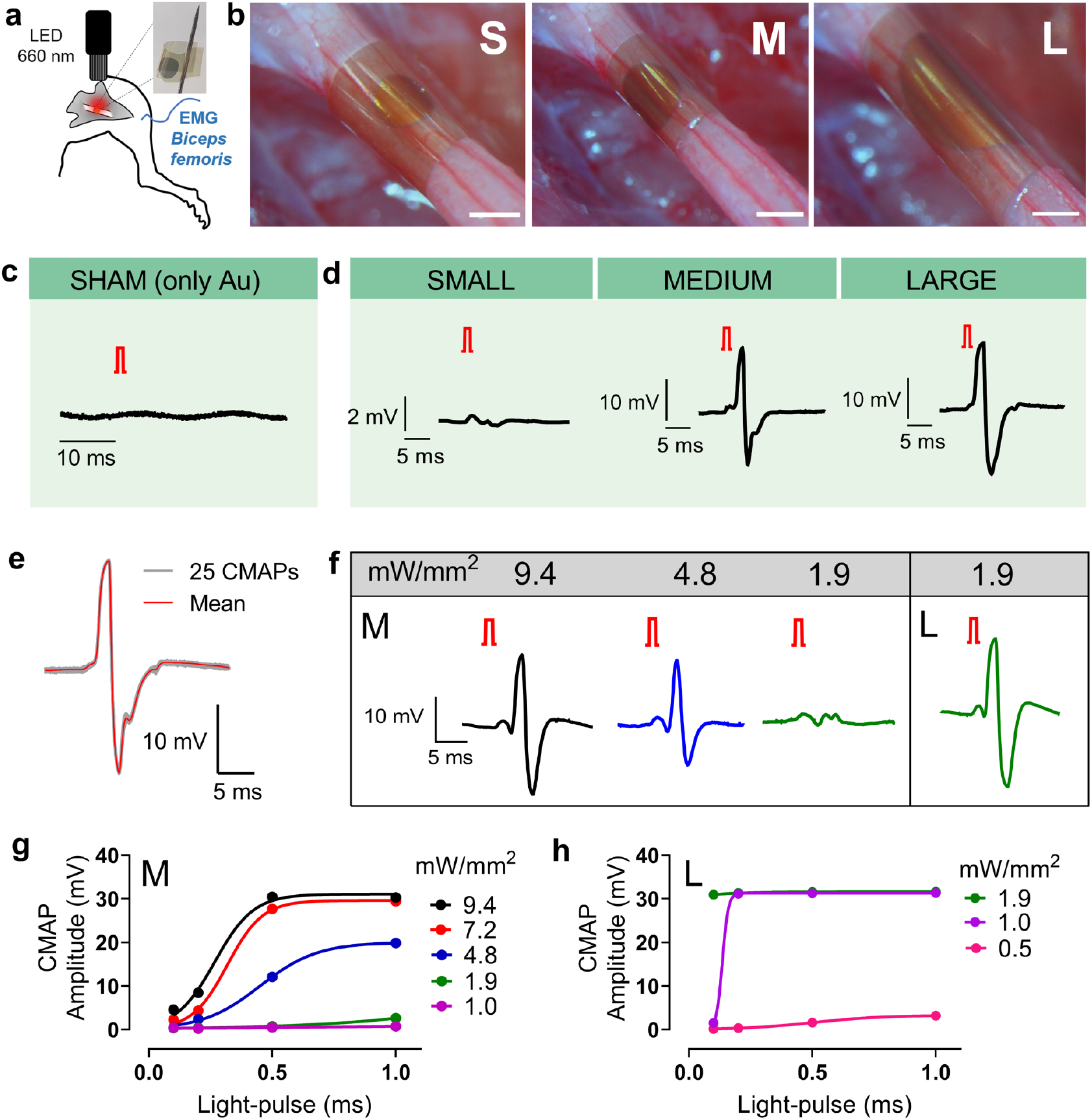
Acute sciatic nerve photostimulation is precisely controlled by varying OEPC device size, light intensity, and stimulation pulse length. **a,** Schematic of the sciatic nerve stimulation experiment design for acute conditions, with inset photograph showing a free-standing device prior to implantation. **b,** Photographs of S, M, and L devices wrapped around the sciatic nerve. Scale bar 1 mm. **c,** Illumination of a sham device, without the PN pixel (only gold on parylene C) gives no response or artefact. 1-ms light-pulses, 9.4 mW/mm^2^ irradiation. **d,** Averaged evoked CMAPs in the biceps femoris (Rat #1) during 25 repetitive light-pulse stimulation (1 ms pulses, 9.4 mW/mm^2^ irradiation, 3 seconds in-between), for the respectively-sized OEPC devices. **e,** Highly reproducible repeated stimulation can be evidenced when 25 CMAPs are plotted on top of each other (in grey) for biceps femoris after repetitive stimulation with a M-sized OEPC device (Rat #1, 1 ms light pulses, 9.4 mW/mm^2^ irradiation, 3 seconds in-between). The averaged response is shown in red. **f,** Examples of CMAPs evoked at different light intensities, 1 ms pulses on M- and L-sized OEPC devices (Rat #2) **g,** Average biceps femoris CMAP amplitudes versus light-pulse length, at different light intensities. M-sized OEPC device. 25 pulses, 3 seconds in-between, for each condition. **h,** Average biceps femoris CMAP amplitudes versus light-pulse length, at different light intensities. L-sized OEPC device. 25 pulses, 3 seconds in-between, for each condition. CMAP amplitude saturates at lower intensities and pulse times for the L device compared with the M device.

*In vivo* neurostimulation often requires precise control of response timing and amplitude. Using M- and L-sized OEPC devices, we therefore determined the relationship between light intensity and light pulse duration with CMAP responses (Figure 3f-h, Figure S1e-g). For size M devices, light intensities in the range of 4-10 mW/mm^2^ evoked robust CMAP responses, and visible movements in the leg and paw (Figure 3f,g, Figure S1e,f). Light-pulse durations between 200-500 μs were needed to reach a threshold of visible movement. With the L-sized OEPC device, the light intensity and light-pulse duration could be significantly reduced (Figure 4f,h, Supplementary Figure S1e,g). For instance, at 1 mW/mm^2^, using 1-ms light-pulses, the CMAP response already saturated for both muscles (Figure 3h, Figure S1g). Consequently, shorter light-pulse lengths (100 μs) could be used to reach a threshold of visible movement. Therefore, light intensity, light-pulse duration, and PN pixel size interplay to produce a given level of OEPC-mediated neurostimulation.

**Figure 4.**
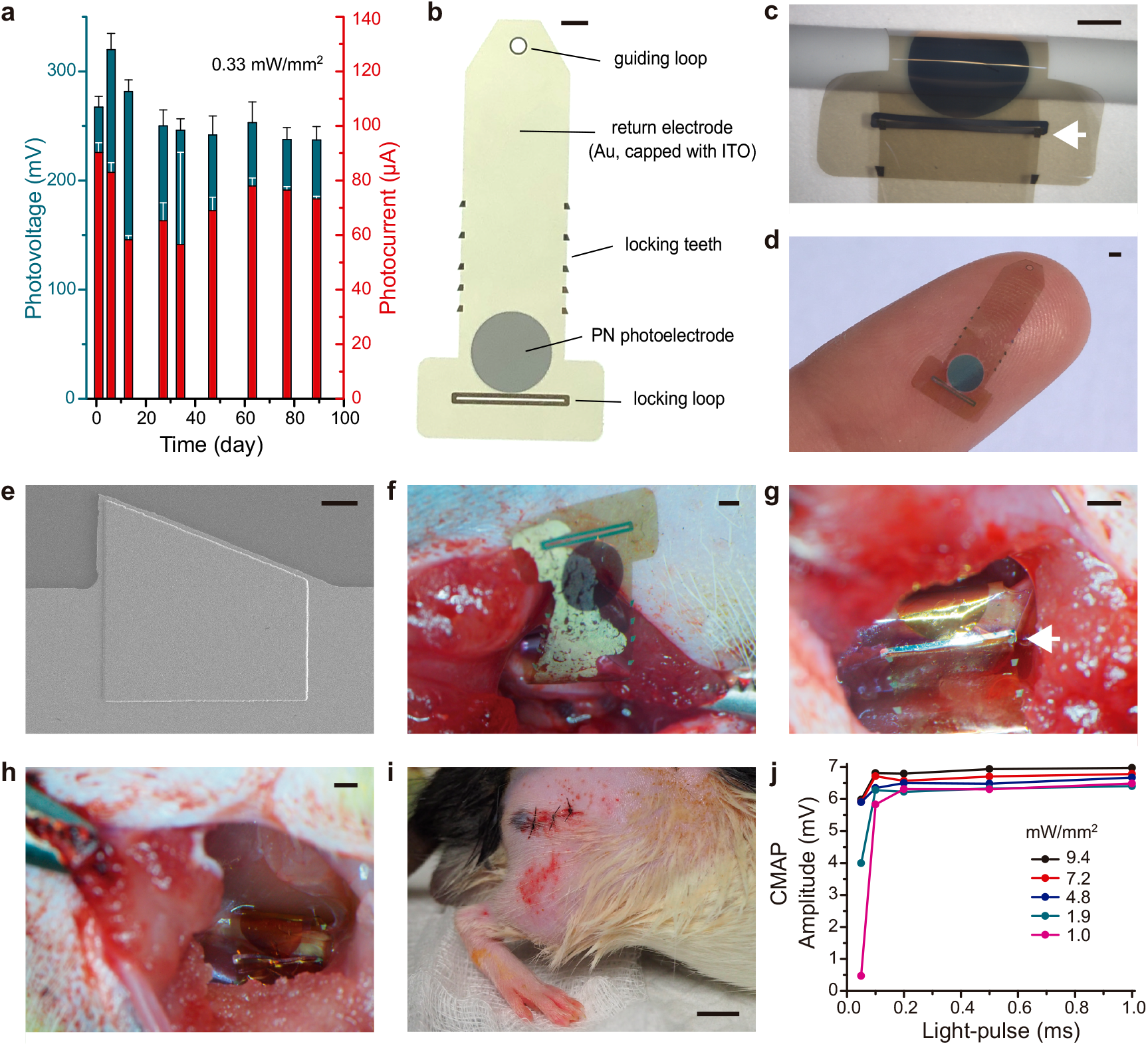
Self-fixating OEPCs are mechanically robust to intra-operative placement and chronic *in vivo* implantation. **a,** Verification of *in vitro* stability: Photovoltage and photocurrent recorded periodically for devices subjected to accelerated aging conditions in 0.1 M KCl solution at 42 °C with light pulsing stress totaling 14 million charge/discharge cycles. n = 4, avg. ± SD. **b,** Photograph of the zip-tie CIP device with labeled components, and **c,** when wrapped and locked around a nerve phantom (1.4 mm diameter cylinder). The white arrow indicates the locking tooth. Scale bars 1 mm. **d,** Photograph of the conformable, ultralight and biocompatible CIP final design; scale bar 1 mm. **e,** SEM micrograph of the parylene-encapsulated aluminum tooth; scale bar 40 μm. **f,** CIP implantation was initiated by inserting the end of the ribbon behind the nerve and tucking it below the nerve; scale bar 1 mm **g,** The PN pixel was positioned adjacent to the external facing surface of the nerve and the end of the ribbon was pulled through the loop to lock the ratchet teeth against the loop edges (white arrow); scale bar 1 mm. **h,** CIP device closed around sciatic nerve, scale bar 1 mm. **i,** Sutured incision after CIP implantation was complete, scale bar 1 cm. **j,** Relationship between light intensity/pulse time and CMAP amplitude for implanted CIP device prior to incision closure.

### Chronically implantable photocapacitor (CIP) development and implementation

Following acute nerve stimulation experiments, we aimed to engineer the OEPC for chronic implantation by developing a more chemically and mechanically stable interface with the nerve. The OEPC relies on an ultrathin Au semitransparent conductive layer on parylene C as the return (+) electrode. However, we found that this Au layer loses conductivity and delaminates when stored in chloride-containing electrolytes for more than two weeks. This likely precludes *in vivo* longer-term use. To stabilize the thin Au, we deposited a 30 nm-thick encapsulating layer of indium tin oxide (ITO), which has excellent adhesion with Au^54^. ITO has a wide electrochemical passivity window^55^, is biocompatible^56,57^, and does not compromise transparency or conductivity. Furthermore, the ITO layer enabled convenient new micropatterning approaches for us, based on the parylene peel-off lithography^58^, at it served as an effective etch-stop for O_2_-reactive ion etching (see *Methods*). To predict *in vivo* stability and functionality of CIP devices, we conducted accelerated aging/stressing tests by immersing devices in 0.1 M KCl solution set to 42 °C, and illuminating them with constant light pulses (2 Hz) delivered through a high-density LED array. Devices were periodically removed and photovoltage and photocurrent were registered using the previously-described EPR technique. No devices failed, and photovoltage/current retained greater than 85% of starting values over the course of 89 days of continuous stressing (corresponding to 14 million charge/discharge cycles; Figure 4a). Thus, the ITO modification of the OEPC gives promising indications of robustness. Because the L-sized OEPCs performed optimally at low-light intensities in acute experiments, this PN pixel size was used for fabrication of CIP devices.

To ensure conformable contact with neural tissue and prevent damage caused by tension or pressure, we had chosen from the start to use ultra-thin parylene C substrates for the OEPC neural interfaces. Parylene C can maintain mechanical contact via capillary forces in the presence of water, but this contact was insufficient to immobilize the device on the nerve for long-term stable stimulation. Therefore, we adapted a parylene zip-tie locking mechanism^59,60^ that allowed the device to form a cuff around the nerve in a fixated position. One end of the parylene C substrate ribbon was narrowed and modified to contain a small guiding loop, and the other end widened to contain a narrow locking loop (Figure 4b-d). When the guiding loop ribbon was passed through the locking loop, locking “teeth” along the lateral borders of the ribbon allowed for sizing of the cuff diameter and prevented slippage of the ribbon ends (Figure 4c). This CIP remained fully conformable, and was sized to accommodate the dimensions of rat sciatic nerve accessible through minimally invasive surgical incision. Evaporated aluminum, which is a relatively malleable metal, (1.5 μm thickness) encapsulated with parylene C, reinforced the loops and teeth (Figure 4e). These aluminum-stiffened “teeth” and loops provide mechanical robustness of the locking mechanism and enable facile manipulation with surgical tools during the implantation procedure.

Implantation was performed by sliding the ribbon of the CIP behind the sciatic nerve (Figure 4f) and pulling the guiding loop through the locking loop with fine forceps (Figure 4g). Once advanced through the locking teeth, the excess parylene C ribbon was trimmed and the sciatic nerve resumed anatomic position relative to the muscles of the hindlimb (Figure 4h). This procedure was straightforward and required 5 minutes of surgical time, fulfilling a key criterion for translation of such devices to practical applications^61^. The incision was then sutured, separating the CIP from the surface of the skin by approximately 10-15 mm of skin, subcutaneous tissue, and muscle (Figure 4i). CMAPs evoked by CIP devices had comparable light intensity-dependent amplitude responses to acutely implanted OEPCs, confirming that design modifications to enable chronic implantation did not adversely affect functionality (Figure 4j).

We implanted CIP devices into a cohort of six rats and monitored functionality at regular intervals over the course of three months. To minimize animal discomfort, we recorded CMAPs through gel-based cutaneous electrodes (Figure 5a). Light stimulation from outside the body (using a 638 nm diode laser with a 2 mm ø illumination spot, 700 mW max as shown in Figure 1d-f) elicited robust and repeatable CMAPs in all rats, some of which were associated with large-amplitude muscular twitches (Figure 5b, Supplementary Video 2).

**Figure 5.**
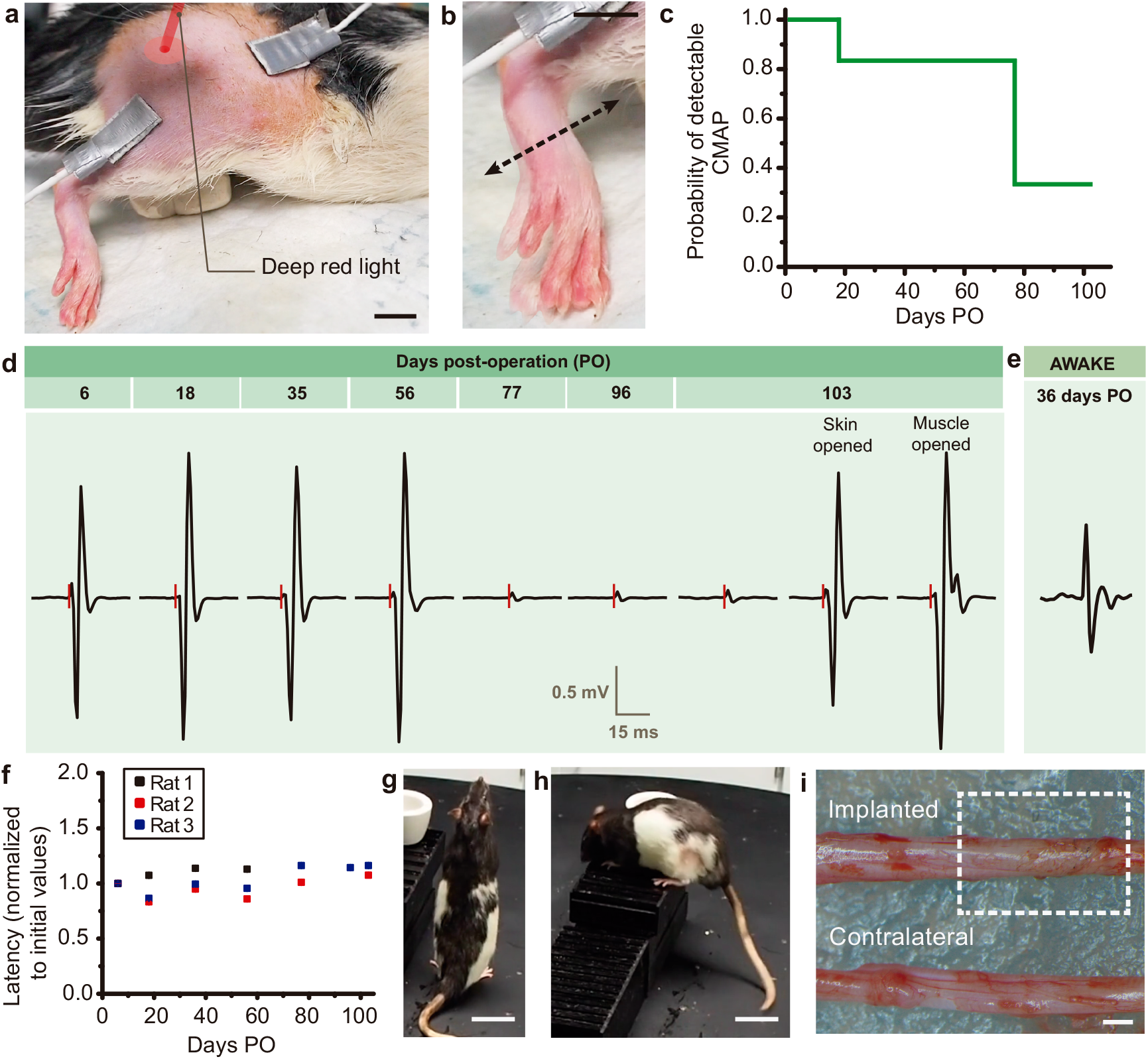
OEPCs permit chronic, non-invasive *in vivo* sciatic nerve stimulation. **a,** Healed implanted area after post-surgical recovery with surface electrodes for EMG measurement attached; scale bar 1 cm. **b,** Photoinduced stimulation with 200-1000 μs at 700 mW induced observable muscular twitches; scale bar 1 cm; see Supplementary Video 2. **c,** Survival curve demonstrating the persistence of detectable CMAPs over the implantation period for all rats (n = 6). **d,** Representative averaged CMAPs over 103 days of chronic implantation. CMAP amplitude abruptly decreased by 77 days post-operatively (PO), but the device was revealed to be functional upon increased light delivery accomplished by opening the overlying tissue. The red dashes indicate light stimulation pulses **e,** Representative photoinduced CMAPs recorded in an awake rat. **f,** Stable average latency between light pulse and CMAP peak over initial implantation period for the three rats with functional devices exceeding 60 days. **g-h**, All rats (n = 6) exhibited normal behavior without noticeable hindlimb-related movement impairments, including standing on hindlimbs, climbing or running (see Supplementary Video 4); scale bar 5 cm. **i,** Implanted sciatic nerve area did not show gross pathological changes compared to the unimplanted contralateral sciatic nerve; scale bar 1 mm.

CIP devices generated detectable CMAPs in all rats for 20 days, and the majority were functional for longer than 60 days (Figure 5c). In two rats, devices remained functional at timepoints greater than 100 days post-operatively, and CMAPs were obtainable in both the anesthetized and awake states (Figure 5d,e). Furthermore, when CIPs were functional, latency of CMAP onset from application of light stimulus was relatively constant for light stimuli of variable intensity and duration (Figure 5f), indicating stable nerve conduction velocity at values expected for the intact rat sciatic nerve^62^. We observed that CMAP amplitude was typically stable for a prolonged time period without decay, but could experience abrupt decrement (Figure 5d, left to middle) that was associated with slightly increased CMAP latency (Supplementary Figure S2). However, CIP devices remained capable of eliciting CMAPs with similar amplitude and latency to those obtained shortly after implantation when light intensity was increased by eliminating tissue attenuation (Figure 5d, right, Supplementary Video 3, Supplementary Figure S2). Given these results, we hypothesize that the alteration in CMAP properties during the implantation period was related to generation of a submaximal neural excitatory pulse due to decreased CIP stimulation performance (*i.e.* device degradation) rather than compromise of the nerve^63^. Post-mortem explantation of CIP devices revealed partial delamination of organic PN material from the parylene C substrate (Supplementary Figure S3). This observation of device degradation is in contrast to the results of the accelerated ageing tests *in vitro* shown in Figure 4a. These findings highlight the importance of performing chronic *in vivo* experimentation in order to subject devices to the full range of biochemical and mechanical stressors present in a freely moving organism.

None of the rats demonstrated motor deficit at any time post-implantation. Each rat was capable of running, standing on hindlimbs, and climbing (Figure 5g-h, Supplementary Video 4), suggesting that CIP implantation does not impair motor function. Consistent with this notion, there were no gross pathological differences between the implanted and contralateral sciatic nerve on post-mortem examination (Figure 5i).

## Discussion

We have demonstrated that ultrathin organic photocapacitors can be microfabricated into conformable devices that are capable of generating sufficient electrical charge to modulate neural tissue *in vivo*. Device-mediated neuromodulation is accomplished via noninvasive, tissue-penetrating deep red-light. Chronically implanted photocapacitors exhibit physiologic stability and functional stimulation of a peripheral nerve over months in a freely-moving animal and do not incur motor deficit. In contrast to many other wireless neuromodulation devices, such as electromagnetic induction or ultrasound-based transducers, photocapacitors are microfabricated in a thin film configuration, resulting in a minimally invasive interface with tissue.

Photocapacitors offer several advantages compared to other stimulation modalities. Conventional electrical stimulation is accomplished by wired leads connected to an implanted power source, a configuration that is a common cause of device complications^64^. We show that photocapacitors are capable of chronic electrical stimulation by directly converting light impulses into charge-balanced biphasic electrical signals, which is considered favorable for safe long-term stimulation^51^. Furthermore, the charge generated, and thus the neural response elicited, are directly related to the strength and duration of the light pulse. These features enable precise temporal and amplitude control of stimulation patterns. Because photocapacitors are microfabricated, they are inherently customizable. The size, configuration, and location of PN junctions can be modified within a variable shape and size of ribbon substrate, permitting application to nerves of different diameters, as well as other types of neural tissue. Our flexible locking mechanism minimizes risk of tissue damage while maintaining steady device position *in vivo*. In addition, no power source apart from the light pulse is necessary to operate the device, eliminating risks associated with implanted power hardware^65^. Photovoltaics based on the same active components as used in CIPs could also conceivably be used to non-invasively power other electrical components and enable information transmission via narrow-band LEDs and photodiodes.

To facilitate translation of photocapacitor devices to clinical applications, sustained performance should be demonstrated over prolonged time periods. Current CIP devices functioned for months in a freely moving rodent, establishing potential feasibility. There are three clear areas for optimization of CIPs: device stability, efficiency, and higher light sensitivity at longer wavelengths. Device longevity could be improved by encapsulation of the PN pigment with a conductive layer that prevents exposure of the internal device layers to electrolyte without decreasing electrical performance. The second parameter to optimize is light-to-charge efficiency of the devices, which would allow for operation in deeper tissues and increase the variety of targetable neural structures. Alternatively, it is possible to microfabricate conformable circuits that transmit electrical charge to deeper structures while maintaining the photoactive pigments closer to the external tissue interface for effective light activation. However, much can be gained by tuning the stimulation wavelength. According to our MC model, a wavelength of 700 nm would be optimal in terms of transmission, and would nearly double the possible implantation depth (Supplementary Figure S4). On the other hand, tuning photocapacitor devices to respond even further to the red, 800-900 nm, could also be advantageous for comfort of human subjects, as at these wavelengths tissue transparency is sufficient for device operation, but human vision is no longer sensitive^36^.

Electrical neurostimulation is employed not only to assay neural function in experimental paradigms^66^, but is an efficacious and well-tolerated therapy for multiple neurologic disorders, from chronic pain to epilepsy^67^. CIPs can facilitate testing of such neuromodulatory protocols in small animal models by minimizing device footprint and allowing for full device implantation without any tissue traversing elements, features that have been demonstrated to improve experimental outcomes^68,69^. The unique features of CIPs also advance the potential for translation to bioelectronic devices that require safe, long-term neurostimulation to treat pain and enable motor rehabilitation in humans.

## Methods

### OEPC device fabrication

H_2_Pc, (Phthalocyanine, Alfa Aesar) and PTCDI (*N,N′*-dimethyl- 3,4,9,10-perylenetetracarboxylic diimide, BASF) were first purified by threefold temperature-gradient sublimation. 4-inch soda lime glass wafers (University Wafer, 550 μm thick) were cleaned in a circulating 2% solution of Hellmanex III detergent heated to 50°C for 30 min followed by a high-pressure rinse with acetone and deionized water (DI). The wafers were then treated with O_2_ plasma (Diener electronic GmbH, 200 W, 20 min). Immediately after, a 5 μm-thick parylene C layer was deposited via chemical vapor deposition (Diener electronic GmbH). The parylene C was then patterned by lithography and etching to produce 4 ×15 mm ribbons. This was done as follows: An aluminum reactive ion etching (RIE) hard mask was deposited through a stainless steel shadow mask onto the parylene C wafer. The 80 nm layer of Al was evaporated in a PVD chamber in a vacuum of < 2×10^−6^ Torr using a rate of 5-15 Å/s. Parylene C was removed by RIE (200W, O_2_ 100 sccm). Finally, the Al mask was etched using a commercial wet etch solution. The parylene surface was then activated with O_2_ plasma (50 W, 2 min), followed by vapor-phase treatment with 3-(mercaptopropyl)trimethoxysilane, MPTMS, by placing the samples in an MPTMS-vapor saturated chamber heated to 90 °C for 1 h. MPTMS treatment enhanced the adhesion of Au to the parylene C substrate. Next, a 10 nm-thick film of Au was thermally evaporated over the whole wafer in a vacuum of < 2×10^−6^ Torr using a rate of 3-5 Å/s. The organic pigment PN pixels were formed by thermal evaporation through a shadow mask at a base pressure of < 2 × 10^−6^ Torr using a rate of 0.1-0.5 nm/s. 30 nm of P-type H_2_Pc and 30 nm of N-type PTCDI were successively deposited resulting in 60 nm total thickness of the organic layers (PN). It should be noted that efforts were made to produce semitransparent contacts from other materials, such as ITO. However, due to poor adhesion to the parylene substrate, approaches with ITO alone proved unsuccessful.

### Optical tissue penetration modeling

Monte-Carlo light propagation simulation was conducted using the CUDAMCML software^47^, a gpu-accelerated version of the well-established MCML software^46^. The tissue model consisted of three layers, a 1 mm layer of skin followed by 3 mm of subcutaneous adipose tissue and a 50 mm thick layer of muscle. Rat tissue optical parameters for all the wavelengths from Bashatkov and coworkers^48^ were used. Light penetration was evaluated on a 0.1 mm vertical and radial mesh. 5 billion photons were used for each run of the simulation. The output files were convolved by the CONV software (a part of the MCML distribution), assuming a 2 mm FWHM diameter Gaussian light beam normalized to 700 mW of total power.

### Photoelectrochemical characterization

Measurements of photovoltage and charging current of OEPC devices was performed according to previously described methods^43^. Briefly, the backside Au of the OEPC was contacted with a probe electrode connected to the positive terminal of an oscilloscope. Meanwhile, the negative terminal was connected to an Ag/AgCl electrode in 0.1 M KCl electrolyte, making contact to the top of the organic layer of the OEPC device. The droplet contact area was 0.126 cm^2^. Pulsed illumination was provided by a 638 nm laser diode or a 660 nm high-power LED at various light intensities. Light intensity was verified using a calibrated Thorlabs Si PIN diode (Thorlabs SM1PD1B). The transient voltage change (V_t_) was measured when the OEPC device was immersed in 0.1 M KCl, using a GeneClamp 500B amplifier (Axon Instruments) and a Digidata™ 1440A converter (Molecular Devices), as described previously^43^.

### Acute sciatic nerve stimulation

All animal experiments were approved by the institutional *Animal Care and Use Committee* of Columbia University Irving Medical Center. The implantations were carried out in Long Evans rats (200-250g at the time of implantation) that had no previous experimentation. Animals were housed in pairs, in a regular 12h/12h light/dark cycle and had access to food and water *ad libitum*.

Rats were anesthetized using 3% isoflurane and surgical site was shaved, disinfected and local analgesia was applied. A longitudinal incision (~1cm) along femoral axis was performed and the sciatic nerve was visualized. The connective tissue surrounding the nerve was minimally dissected to release and expose a longitudinal nerve section approximately 4mm long. OEPC devices were wrapped around the nerve, facing the PN pixel to the nerve surface, and fixed through adhesion of the parylene ribbon ends via capillary forces. 3 PN pixel diameter sizes (1, 1.4 and 3 mm) were tested. Tungsten microwires (diameter 50 μm) were placed in biceps femoris and plantar muscles, each providing a separate recording channel referenced to a microwire in the paraspinal subcutaneous tissue. CMAPs were recorded using a custom designed board based on an AD8237 differential amplifier chip and an RHD2000 board (Intan Technologies) for digitization. Illumination was provided by a 660 nm high-power LED (M660L4) with a collimator lens (SM2F32-B) controlled by a ThorLabs DC2200 High-Power LED. A minimum of 25 light pulses (3 s in between) was used for each light intensity/duration condition tested. Light intensity was verified with a Thorlabs SM1PD1B Si PIN diode. At the conclusion of the intra-operative recording session, the rats were euthanized. 5 rats were used for acute sciatic nerve experiments that aimed to characterize the performance of OEPC devices in regards of PN pixel size, light intensity, and light pulse duration.

### Chronically implantable device fabrication

Clean 4” wafers were coated with 2.2 μm parylene C. The surface was then activated by an oxygen plasma treatment (50W, 120s) followed by physical vapor deposition of 1.5-2 μm thick layer of aluminum (vacuum <1 × 10^−5^ Torr, 20-30 nm/s) acting as mechanical support for the locking mechanism structures.

S1818 photoresist was spin coated on the substrates, exposed with MA6/BA6 Süss Mask Aligner and developed in MF-319. The aluminum layer was patterned using a commercial acidic etchant. The resist was stripped in acetone, followed by isopropanol and deionized water (DI). Propylene carbonate was then spin coated on the wafers at 2000 rpm and baked at 60°C for 30s to act as an adhesion promoter^70^ for the next 2.2 μm thick encapsulating layer of parylene C. The surface was then exposed to oxygen plasma (50W, 120s) and vapor treated with MPTMS in a closed chamber at 80°C for 2h. The wafers were then washed with acetone, isopropanol and DI. Next, 10 nm of gold and a 30 nm of indium tin oxide (ITO) were sputtered onto the substrates. The sputtering target was In_2_O_3_/SnO_2_ 90/10 wt%, 99.99% pure (Lesker). Gold was deposited at 100 W, 3.4 mbar, 100% Ar, DC; ITO at 32 W, 4.5 mbar, 97% Ar, 3% O_2_, RF. The outline of the implants was patterned using S1818 photoresist, MF-319 developer, ITO etchant (concentrated HCl), Au etchant (KI/I_2_) and finally by RIE (200W, O_2_ 100 sccm). The wafer was washed in acetone, isopropanol and DI. A thin layer of 2% cleaning agent Micro90 was spin coated at 1000 rpm to act as an anti-adhesive layer before the next 2.2 μm thick sacrificial layer of parylene C. The openings for the photopixels were patterned by AZ 10XT resist, AZ developer and RIE (200W, O_2_ 100 sccm). The wafers were sequentially washed with acetone, isopropanol and DI. Then the protective layer of ITO was etched with concentrated HCl for a few seconds, followed by a quick wash in DI. Next, the organic pigments H_2_Pc and PTCDI were evaporated from resistively heated crucibles at 1 × 10^−6^ Torr at rates of 1-6 Å/s to produce a 30/30 nm PN junction. Finally, the sacrificial parylene layer was peeled off to yield the patterned device. The wafer was rinsed with DI and tested using the EPR setup described above. In addition, control devices were subjected to an accelerated aging and light-stressing test according to the method described previously^44^.

### Chronic implantation test on the sciatic nerve

Rats were anesthetized and the sciatic nerve was exposed as previously described for acute stimulation. The CIP ribbon was passed behind the exposed sciatic nerve and the end of the ribbon was inserted through the ribbon loop using forceps. The PN pixel was placed facing the nerve surface, and the ribbon was pulled until the teeth passed through the ribbon loop, firmly closing the zip-tie mechanism. The ribbon was adjusted to fit snugly around the nerve without applying compressive force. The excess ribbon (~5mm) was then cut. Sutures were used to close the incision. After the operative procedure was complete, anesthesia was removed and the rats were allowed to recover from surgery. Triple antibiotic ointment and injectable analgesia were applied during the post-surgical recovery period.

The CMAP recording sessions were performed at 6, 18, 35, 56, 77, 96 and 103 days post-operatively (PO). Rats were anesthetized (3% isoflurane) and the implanted site was shaved to facilitate the attachment of EMG electrodes to the skin. Because multiple recording sessions were planned for each rat, we performed non-invasive CMAP monitoring to prevent ongoing disruption of muscle tissue. Gel electrodes (14 × 9 mm, Acuzone) were attached to the skin using Elefix conductive paste (Nihon Kohden). Electrodes were placed in gastrocnemius and vastus lateralis (reference electrode) muscles, maintaining a consistent inter-electrode distance across sessions. Photostimulation was induced by a 638 nm laser diode with a maximum output power of 700 – 1200 mW driven by ThorLabs DC2200 High-Power LED controller. 250 light pulses (3s interpulse interval) with an intensity spanning 700 mW to 7 mW and duration from 1 ms to 0.05 ms were used. CMAP signals were recorded as during the acute stimulation sessions. Video recordings of muscular twitches were performed.

Motor performance of the rats was evaluated in the immediate post-operative period and tested on an open field maze with horizontal and vertical obstacles at 53-54 days post-operatively. Walking and running gait, as well as ability to stand on hindlimbs and climb were observed.

In the subset of rats that showed responses after 103 days PO, the CMAPs were additionally recorded after creating an incision over the implantation site and opening the skin. Subsequently, photoinduced stimulation was repeated as per the above parameters after exposing the device to quantify device performance under maximal light intensity conditions.

At the end of the implantation period, the rats were euthanized, and the implanted sciatic nerve section was harvested and dissected. Additionally, the contralateral sciatic nerve anatomically corresponding to the implanted region was harvested for comparison. Gross pathological examination was performed for all harvested nerve segments. A total of 6 rats were used for chronic sciatic nerve experiments.

## Author Contributions

M.S.E.; M.J.; J.F.L.; and L.M. contributed equally to this manuscript. M.S.E.; M.J.; and L.M. carried out the photoelectrochemical characterizations. M.S.E.; M.J., V.Đ.; D.K.; J.G. performed acute experiments and analyzed data. L.M. fabricated devices for acute experiments. M.J. fabricated devices for chronic experiments. V.Đ. wrote programs for data acquisition, processing, and all modeling. Z.Z. designed and developed the electrophysiological acquisition hardware. J.F.L; D.K.; and J.G. performed the rodent surgeries. Chronic data was collected and analyzed by M.J.; J.F.L.; Z.Z.; J.G.; and E.D.G. The project was led and supervised by M.B.; D.K., V.Đ.; J.G.; E.D.G. The manuscript was written with input from all coauthors.

## Acknowledgements

The authors gratefully acknowledge financial support from the Knut and Alice Wallenberg Foundation within the framework of the Wallenberg Centre for Molecular Medicine at Linköping University, the Swedish Research Council (Vetenskapsrådet, 2018-04505), and the Swedish Foundation for Strategic Research (SSF). This work has been supported in part by Croatian Science Foundation under the project UIP-2019-04-1753. This work was also supported by Columbia University, School of Engineering and Applied Science; as well as Columbia University Medical Center, Department of Neurology and Institute for Genomic Medicine.

## Data Availability

All data supporting the results drawn from experiments can be found in the paper and supplementary information. Raw data used for the plots found in Figures 2-5, and S1-S2 are available from the authors upon request.

## Supplementary information

**Figure S1.**
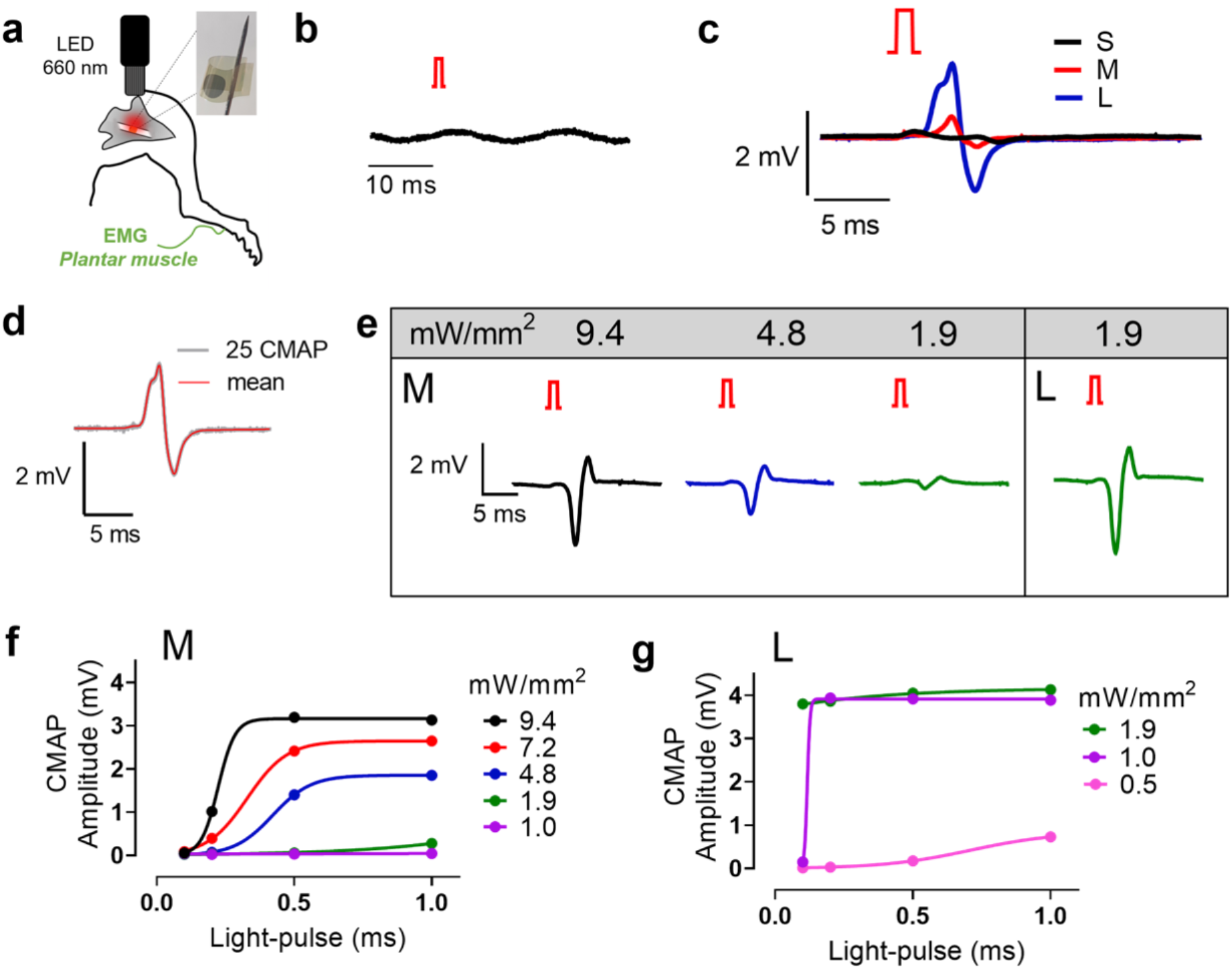
Acute sciatic nerve photostimulation registered in the Plantar muscle. **a,** Schematic of the sciatic nerve stimulation experiment design for acute conditions, with inset photograph showing a free-standing device prior to implantation and EMG electrode inserted into the plantar muscle **b,** Illumination of a sham device, without the PN pixel (only gold on parylene C) gives no response or artefact. 1-ms light-pulses, 8.1 mW/mm^2^ irradiation. **c,** Averaged evoked CMAPs in the plantar muscle (Rat #1) during 25 repetitive light-pulse stimulation (1 ms, 9.4 mW/mm^2^ irradiation, 3 seconds in-between), for the respectively-sized OEPC devices. **d,** Highly reproducible repeated stimulation can be evidenced when 25 CMAPs are plotted on top of each other (in grey) for biceps femoris after repetitive stimulation with a M-sized OEPC device (Rat #1). 1-ms light-pulses, 9.4 mW/mm^2^ irradiation, 3 seconds in-between. The averaged response is shown in red. **e,** Examples of CMAPs evoked at different light intensities, 1 ms pulses on M- and L-sized OEPC devices (Rat #2) **f,** Average plantar muscle CMAP amplitudes versus light-pulse length, at different light intensities. M-sized OEPC device. 25 pulses, 3 seconds in-between, for each condition. **g,** Average plantar muscle CMAP amplitudes versus light-pulse length, at different light intensities. L-sized OEPC device. 25 pulses, 3 seconds in-between, for each condition. CMAP amplitude saturates at lower intensities and pulse times for the L device compared with the M device.

**Figure S2.**
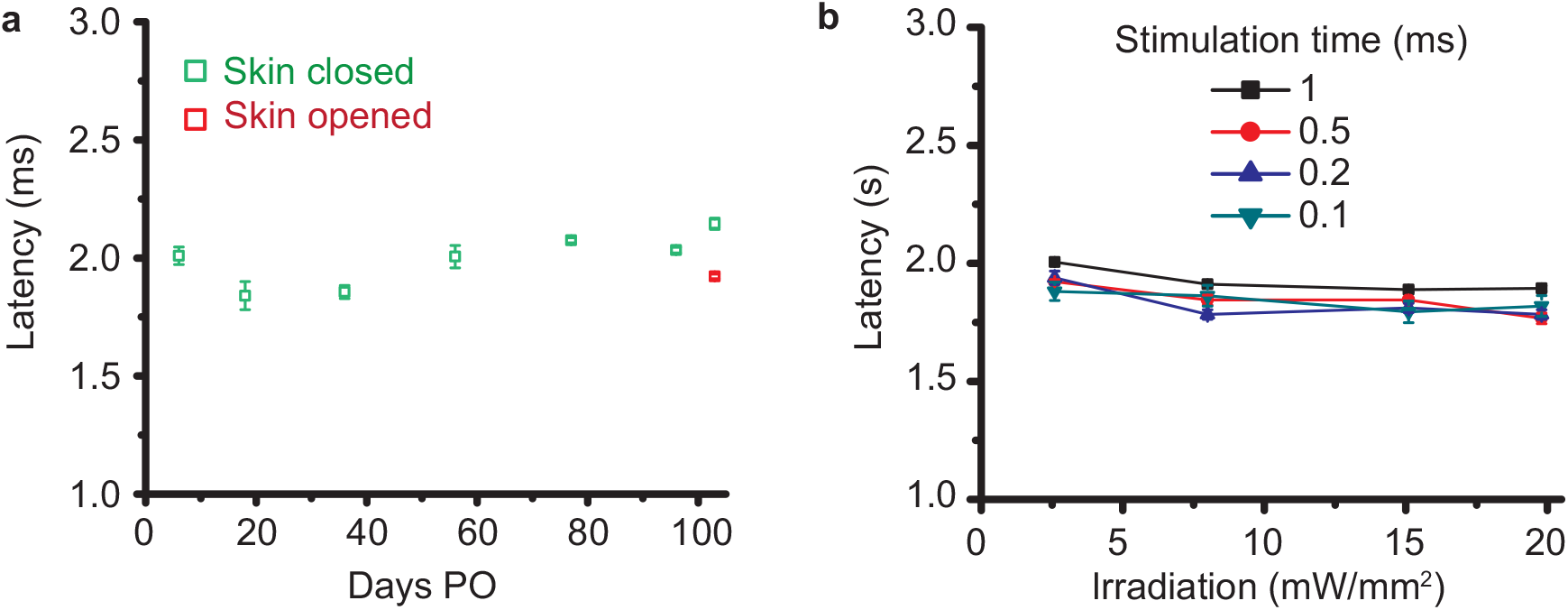
CMAP latency depends on stimulation strength. **a**, CMAP latency over implantation period for a sample rat. Note that although CMAP latency increases after 60 days (green data points), this change is reversible by increasing light intensity through opening of skin superficial to CIP implantation site (red data point). This indicates that the apparent increase in latency is due to drop in device performance rather than detriment to the nerve. **b,** CMAP latency is reduced by increasing light intensity, further confirming that at long implantation times, device performance deterioration is connected with latency rise.

**Figure S3.**
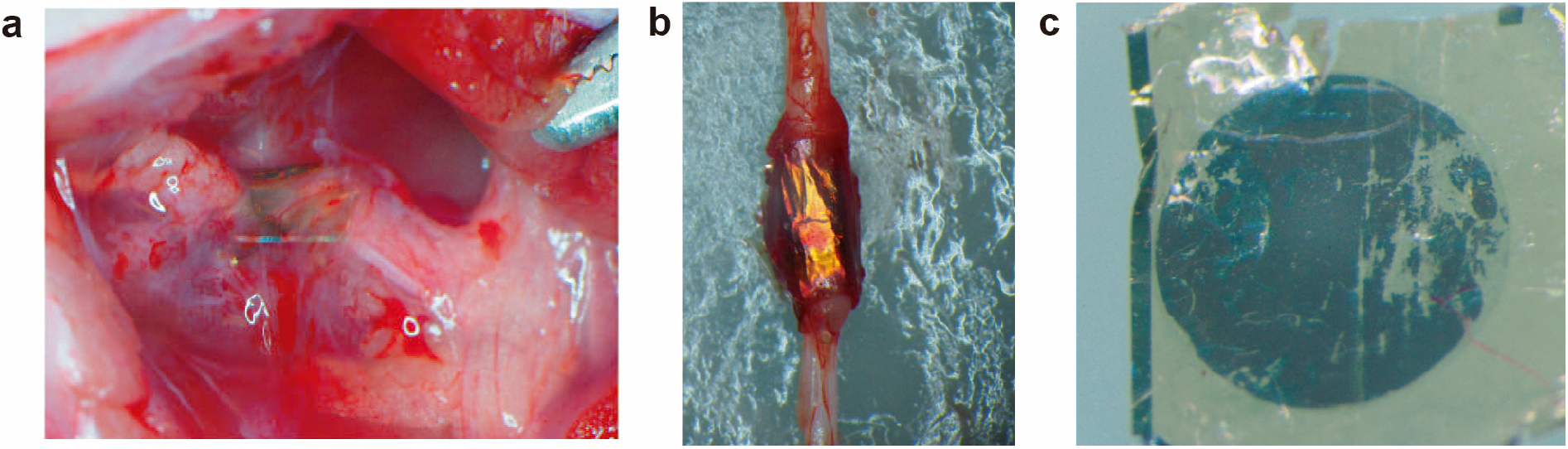
CIP explantation. **a,** CIP affixed to nerve *in vivo* after 103 days post-implantation. **b,** The CIP remained wrapped around explanted sciatic nerve. **c,** Micrograph of the organic PN pigment layers from the explanted CIP device showing focal points of delamination of the organic semiconductor layers. The ITO-capped Au remains visually intact.

**Figure S4.**
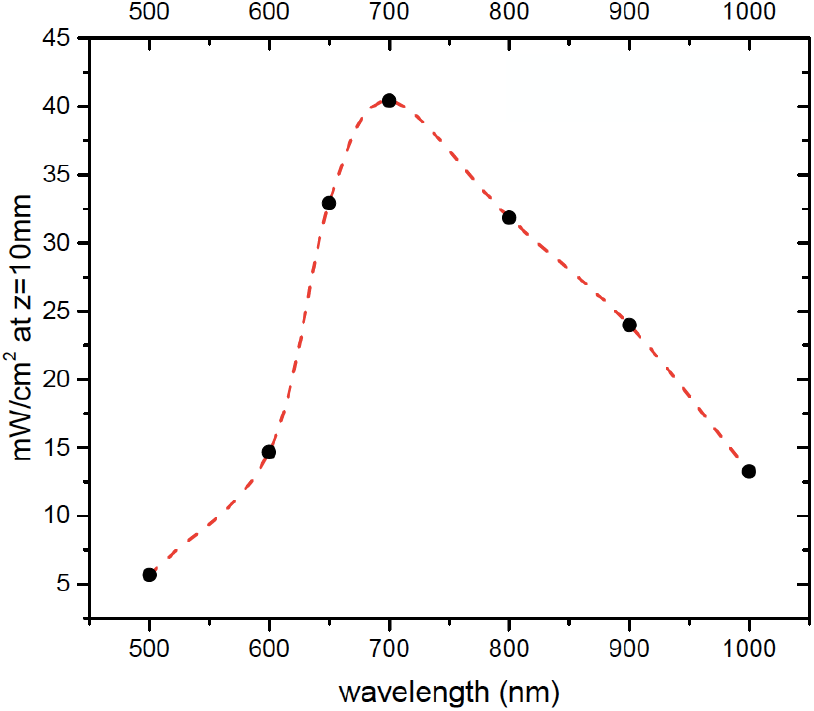
Wavelength dependence of optical power transmission through tissue. Intensity at a fixed depth of z=10 mm below the skin, calculated as a function of beam wavelength. A window between 650 – 800 nm affords the most optimal transmissivity through skin/fat/muscle tissue. The dark circles are points calculated based on available data parameters, while the red dashed line is a guide for the eye.

